# Shugoshin protects centromere pairing and promotes segregation of non-exchange partner chromosomes in meiosis

**DOI:** 10.1101/263384

**Authors:** Luciana Previato de Almeida, Jared M. Evatt, Hoa H. Chuong, Emily L. Kurdzo, Craig A. Eyster, Mara N. Gladstone, Laura Gómez-H, Elena Llano, Régis Meyer, Alberto M. Pendas, Roberto J. Pezza, Dean S. Dawson

## Abstract

Faithful chromosome segregation during meiosis I depends upon the formation of connections between homologous chromosomes. Crossovers between homologs connect the partners allowing them to attach to the meiotic spindle as a unit, such that they migrate away from one another at anaphase I. Homologous partners also become connected by pairing of their centromeres in meiotic prophase. This centromere pairing can promote proper segregation at anaphase I of partners that have failed to become joined by a crossover. Centromere pairing is mediated by synaptonemal complex (SC) proteins that persist at the centromere when the SC disassembles. Here, using mouse spermatocyte and yeast model systems, we tested the role of shugoshin in promoting meiotic centromere pairing by protecting centromeric synaptonemal components from disassembly. The results show that shugoshin protects centromeric SC in meiotic prophase and, in anaphase, promotes the proper segregation of partner chromosomes that are not linked by a crossover.

**SIGNIFICANCE:** Meiotic crossovers form a connection between homologous chromosomes that allows them to attach to the spindle as a single unit in meiosis I. In humans, failures in this process are a leading cause of aneuploidy. A recently described process, called centromere pairing, can also help connect meiotic chromosome partners in meiosis. Homologous chromosomes become tightly joined by a structure called the synaptonemal complex (SC) in meiotic prophase. After the SC disassembles, persisting SC proteins at the centromeres mediate their pairing. Here, studies in mouse spermatocytes and yeast are used to show that the shugoshin protein helps SC components persist at centromeres and helps centromere pairing promote the proper segregation of yeast chromosomes that fail to become tethered by crossovers.

## INTRODUCTION

Faithful chromosome segregation during meiosis depends upon the formation of connections between homologous chromosome pairs. Crossovers, also called exchanges, are the basis these connections. Chiasmata, the cytological manifestation of crossovers, in conjunction with sister chromatid cohesion distal to the crossover, create a physical link that holds the homologous chromosomes in pairs called bivalents (reviewed in (1)). The linkage allows the bivalent to attach to the meiotic spindle as a single unit, such that at anaphase I, the partners will migrate away from one another to opposite poles of the spindle. In some instances, even in the absence of exchanges, proper meiotic chromosome segregation can be achieved (reviewed in (2)). In yeast and *Drosophila* when a single chromosome pair does not experience an exchange, the pair still segregates correctly in most meioses (3–6). Although the behavior of non-exchange chromosomes has been difficult to study in mammalian models, there are indications that here too, there may be mechanisms beyond crossing-over that help to direct their behavior in meiosis I. For example, in mice, the majority of chromosomes in oocytes from Mlh1 recombination-deficient mutant appeared to be spatially balanced on the spindle (7), and in humans, while smaller chromosomes (21 and 22) fail to experience crossovers in about 5% of meioses (8–10), they are estimated to non-disjoin in less than 1% of meioses (9–11).

In yeast and *Drosophila*, the centromeres of non-exchange partners pair or interact in meiotic prophase (12–15). Similar centromere pairing is also seen in mouse spermatocytes (16, 17). Meiotic centromere pairing (or clustering in *Drosophila* females) is mediated by proteins that are components of the synaptonemal complex (SC) (14–18). The SC zippers the axes of homologous chromosomes along their lengths in mid-meiotic prophase (pachytene) and disassembles in late prophase (diplotene). However, all SC components tested (Zip1 in yeast, its functional homologs SYCP1 in mice, and C(3)G in Drosophila, the mouse SC components SYCE1, SYCE2, SYCE3 and TEX12, and the Drosophila protein Cona) persist at centromeres, holding them together in pairs (yeast and mouse spermatocytes; (14–18) or clusters (*Drosophila* females; (18)). In budding yeast, this centromere pairing is correlated with proper disjunction of the non-exchange pair (14, 15).

Important questions regarding the mechanism and function of centromere pairing remain unanswered. First, how does centromere pairing by SC components in prophase ensure disjunction in anaphase? This is especially curious as in budding yeast and mice, the SC components that are protected at the centromeres in late prophase (Zip1/SCYP1) are greatly reduced or undetectable when the centromeres begin attaching to the microtubules of the spindle (14, 16, 17). Second, what enables SC proteins to persist at the paired centromeres when the SC disassembles?

The persistence of centromeric SC in late prophase, when the chromosomal arm SC disassembles, is reminiscent of the protection of meiotic cohesins at centromeres at the metaphase I to anaphase I transition when arm cohesion is lost (reviewed in (19)). Protection of centromeric cohesin in meiosis I is mediated by shugoshin – a function first revealed by studies of the *mei-S322* gene in *Drosophila* (20, 21) (reviewed in (22)). In yeasts and mouse oocytes, shugoshin has been shown to recruit forms of PP2A phosphatase to centromeres, rendering the centromeric cohesin refractory to cleavage by the protease separase at the metaphase I to anaphase I transition (23–27). In budding and fission yeast, shugoshin acts by protecting the Rec8 component of cohesin from phosphorylation by casein kinase, and also in budding yeast by the Dbf4 kinase (DDK). In other organisms the identities of the kinases that prepare cohesins for separase cleavage have not been determined (28–30) (reviewed in (22)).

Phosphorylation also promotes SC disassembly and degradation, but at the pachytene-to-diplotene transition (reviewed in (31)). Studies in rats and mice have correlated the phosphorylation of SC components (SYCP1, SYCP3, TEX12, and SYCE1) with pachytene exit and SC disassembly (32, 33) and the Polo-like kinase PLK1 localizes to the SC central element in pachytene and can phosphorylate SYCP1 and TEX12 *in vitro* (32). Similarly, in budding yeast, Polo-like kinase (Cdc5) expression is central in promoting SC disassembly (34), but it works in a network with other kinases, namely Dbf-4 kinase and cyclin-dependent kinase (CDK) (32, 35–38). In mice, CDK has also been implicated in promoting SC removal.

The parallels between the protection of cohesins and SC components at the centromeres compelled us to explore whether shugoshin is responsible for protecting centromeric SC from disassembly signals upon pachytene exit. Our cytological experiments with mouse spermatocytes revealed that this is the case, while genetic approaches with budding yeast revealed that shugoshin is necessary for mediating the segregation of non-exchange chromosome pairs that depend upon centromere pairing for their meiotic segregation fidelity.

## RESULTS AND DISCUSSION

### Centromere Pairing Links Homolog Pairs in the Absence of Chiasmata

Cytological analyses of prophase chromosome spreads from crossover competent mice have revealed that about 4-5% of prophase chromosomes appear to be achiasmate (non-exchange), (39–42). Previous studies have shown that in diplotene homologous partners that appear to be achiasmate are aligned along their arms but are distinctly connected at their centromeres by a short block of persisting SC (16, 17). This observation suggested that mechanisms exist to form a tight connection between homologous centromeres without the need for homologous recombination. An alternative explanation is that these apparent non-exchange chromosome pairs are connected by crossovers that don’t yield obvious chiasmata in chromosome spreads, for example crossovers very close to the paired centromeres might be undetectable in chromosome spreads. If centromere pairing is dependent upon the formation of undetectable centromere-proximal crossovers that can’t be visualized in chromosome spreads, then in mutants unable to form crossovers, centromere pairing should be greatly reduced. To test this, we examined centromere pairing in mice with mutations that compromise the two major cross-over formation pathways. In mice, *Hfm1* is essential to the formation of Class I crossovers. This pathway gives rise to approximately 90% of crossovers (Class I crossovers), (43, 44). Mus81 is essential for the efficient formation of rarer Class II crossovers (45). We immuno-stained spermatocyte chromosome spreads for SYCP3 and SYCP1, components of the lateral and transverse filaments of the SC, in spermatocytes from wild-type, *Hfm1*^−/−^ and *Hfm1*^−/−^/*Mus81*^−/−^ mice. SYCP3 immunostaining enabled visualization of chromosome cores and SYCP1 was used to monitor persistence of SC components at paired centromeres after SC disassembly (centromere pairing) (16, 17). In each chromosome spread we counted the number chromosome pairs that exhibited no clear chiasmata, whether or not their centromeres were paired (Fig. 1 A and B). In agreement with previous reports, elimination of the Class I, or Class I and Class II pathways resulted in many more achiasmate chromosomes per cell (Fig, 1 C) (43, 44). If centromere pairing of apparently achiasmate chromosomes in wild-type mice, reported previously (16, 17), depends upon undetectable crossovers, then in the recombination mutants achiasmate chromosomes should have a reduced frequency of centromere pairing. To test this, we measured the distance between the centromeres of achiasmate chromosome partners in *WT*, *Hfm1*^−/−^ and *Hfm1*^−/−^/*Mus81*^−/−^ mice (Fig. 1D). The loss of recombination, even in the double mutants had no apparent effect on centromere pairing efficiency (Fig. 1 D, Supl. Fig. 1). This result strongly suggests that the paired centromeres of achiasmate partners are not being held together by undetectable crossovers. But instead relies upon the persisting SC at the centromeres as described previously (16, 17).

**Figure 1.**
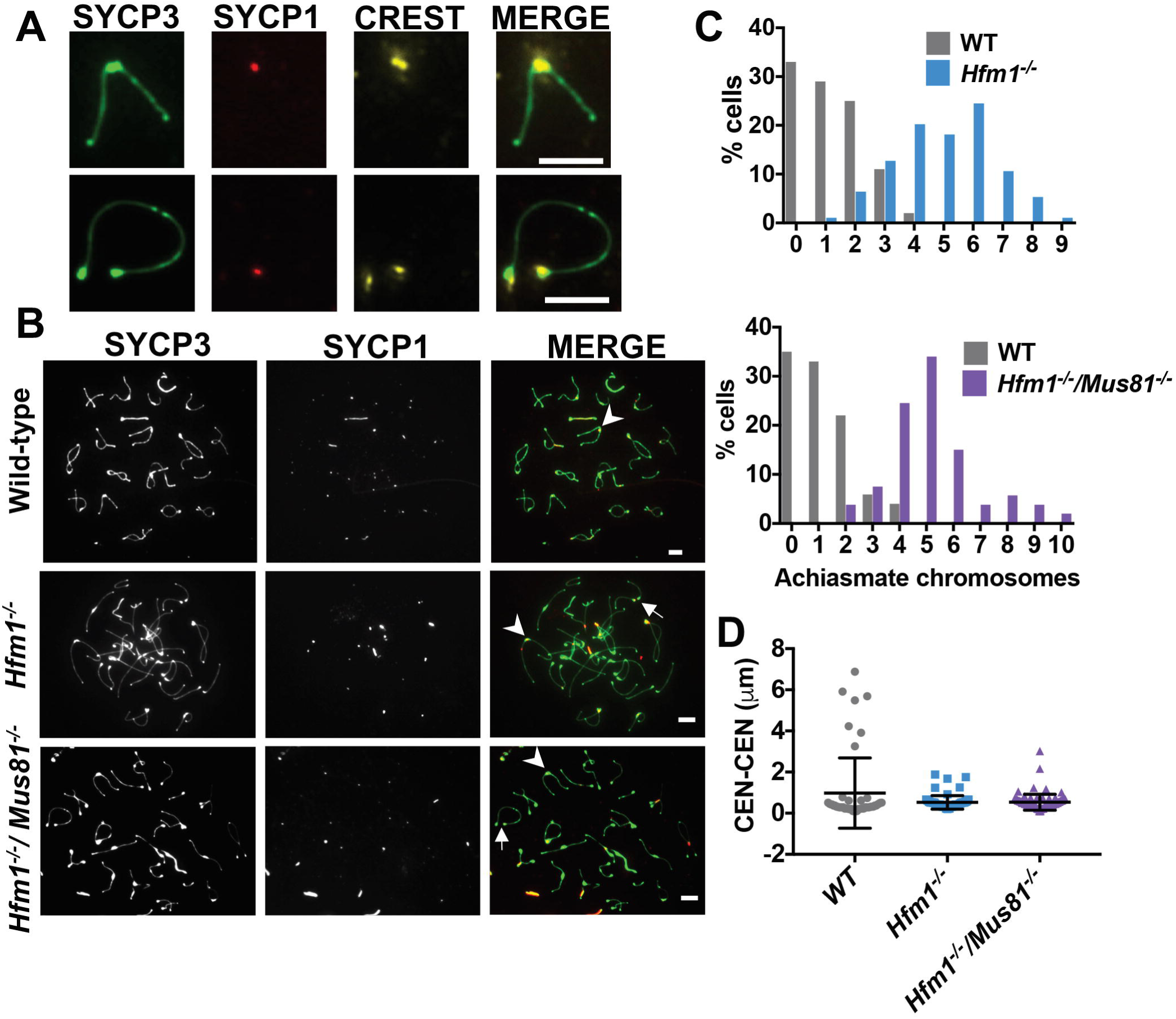
Centromeres pair efficiently in the absence of chiasmata. **A**. Examples of apparently achiasmate partners exhibiting pairing (top panels) or no pairing (lower panels) of their centromeres. Indirect immunofluorescence was used to detect SYCP3 (chromosome axes; green), SYCP1 (synaptonemal complex central element; red), or centromeres (CREST antibody; yellow) in chromosome spreads from mouse spermatocytes. Note that the centromeric end of mouse chromosomes features a bulbous focus of SYCP3 staining and SYCP1 persists at the centromere end after SC disassembly (16, 17). Scale bars equal 5 µm. **B.** Representative mid-diplotene chromosome spreads from wild-type, *Hfm1-/-* and *Hfm1-/- Mus81-/-* mice stained to detect SYCP3 and SYCP1. Arrowheads indicate examples of apparently achiasmate chromosomes with paired centromeres. Arrows indicate examples of apparently achiasmate chromosome partners with unpaired centromeres. **C.** Achiasmate chromosome frequency in spermatocytes from wild-type and recombination deficient mice. Chromosome spreads were scored for the number of clearly achiasmate chromosomes per cell (some chromosomes could not be clearly resolved, thus the graphs under-estimate the achiasmate frequency). Chromosome spreads were from wild-type and mutant littermates. Top graph: WT (gray, n = 100 cells) and Hfm1^−/−^ mutants (blue, n = 94 cells). Bottom graph WT (gray, n = 51 cells) and Hfm1^−/−^ Mus81^−/−^ mutants (purple, n = 53 cells). **D.** The distance between the centromeres was measured for apparently achiasmate partners in the chromosome spreads used for the experiment in panel **C.** WT n = 51 chromosomes; Hfm1^−/−^ n = 69 chromosomes; Hfm1^−/−^ Mus81^−/−^ n = 87 chromosomes.

### Shugoshin 2 protects centromere pairing in mice

Shugoshin 2 (SGO2) localizes to the centromeres of chromosomes in mouse spermatocytes (46). We compared the centromeric localization of SGO2 with the timing of SC protection at centromeres (Fig. 2 A). The axial/lateral element component SYCP3 was used as a marker for the SC. SGO2 is first detected at centromeres in early diplotene cells (Fig. 2 A and B) and remains there thru mid- and late diplotene. This corresponds with the time at which SC components are removed from chromosome arms but not from centromeres (16, 17). Thus, these data are consistent with the model that SGO2 protects the centromeric SC from the disassembly process.

**Figure 2.**
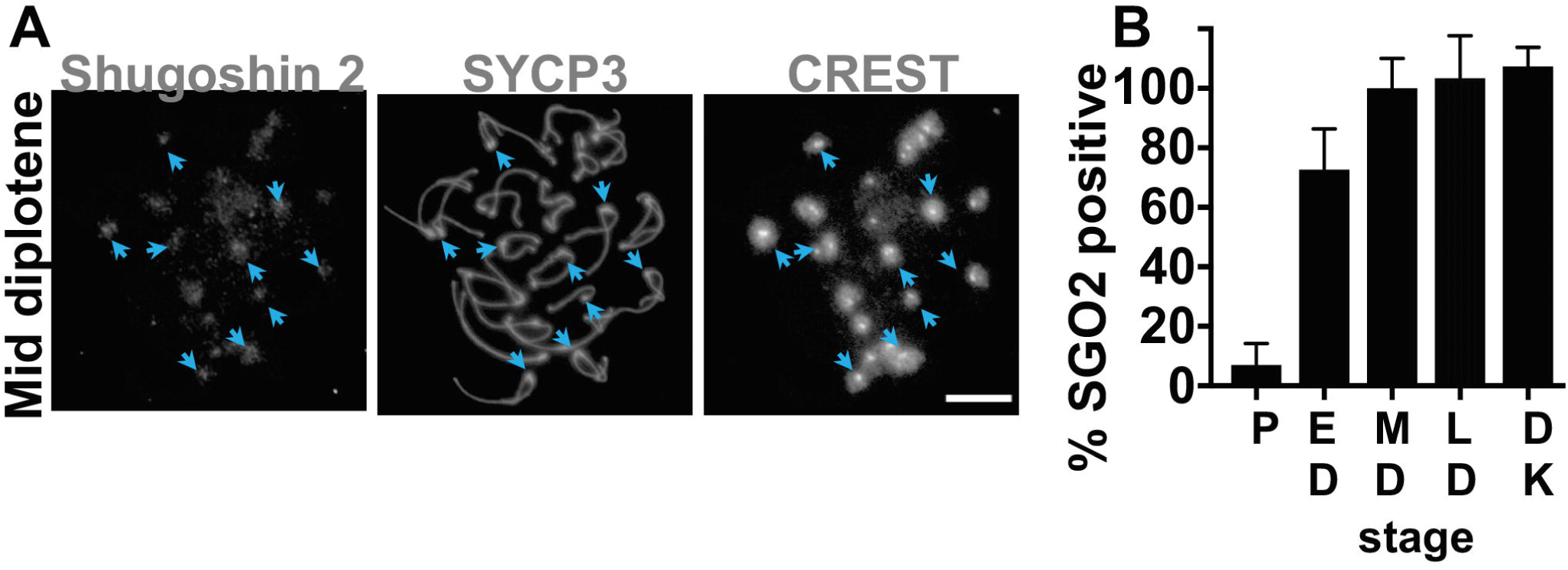
SGO2 co-localizes with persisting synaptonemal complex components at centromeres in prophase of meiosis I. (A) Indirect immunofluorescence was used to evaluate localization of SGO2 on chromosomes at stages of diplotene. Staining with CREST antibody was used to identify centromere regions. Arrowheads indicate examples of paired centromeres. Scale bar represents 5 µm and applies to all images. (B) The average percent of centromeres per spread showing co-localization of SGO2. P = pachytene, ED = early diplotene, MD = mid-diplotene, LD = late diplotene, DK = diakinesis. The stage of each cell was determined by the SC morphology (see Materials and Methods). Error bars indicate standard deviation. A minimum of twenty spreads was scored for each category.

To test whether SGO2 is necessary for the protection of centromeric SC, we monitored the persistence of centromeric SYCP1 in *Sgo2*^−/−^ spermatocytes and wild-type control cells (Fig. 3). In wild-type cells SYCP1 persistence at centromeres mirrors that of other SC components (16, 17). In pachytene cells the SC of the *Sgo2*^−/−^ mutants was indistinguishable from the wild-type control (Fig. 3 A and B). In early diplotene, the SYCP1 signal was visible at nearly all paired centromeres in wild-type chromosome spreads and at nearly 75% of the centromeres in the *Sgo2*^−/−^ mutants (Fig. 3 C). Other SC components, SYCE1, SYCE2 and SIX6OS1, behaved similarly to SYCP1 (Fig. S2). In wild-type spreads the SYCP1 persisted at the centromeres through diplotene (Fig. 3 C). But in *Sgo2*^−/−^ mutants the percent of centromeres with detectable SYCP1 staining was reduced in late diplotene (Fig. 3 A-C). Thus, the absence of SGO2 did not detectably affect SC components in pachytene, but did allow SC components to be lost from centromeres more quickly than in wild-type spermatocytes.

**Figure 3.**
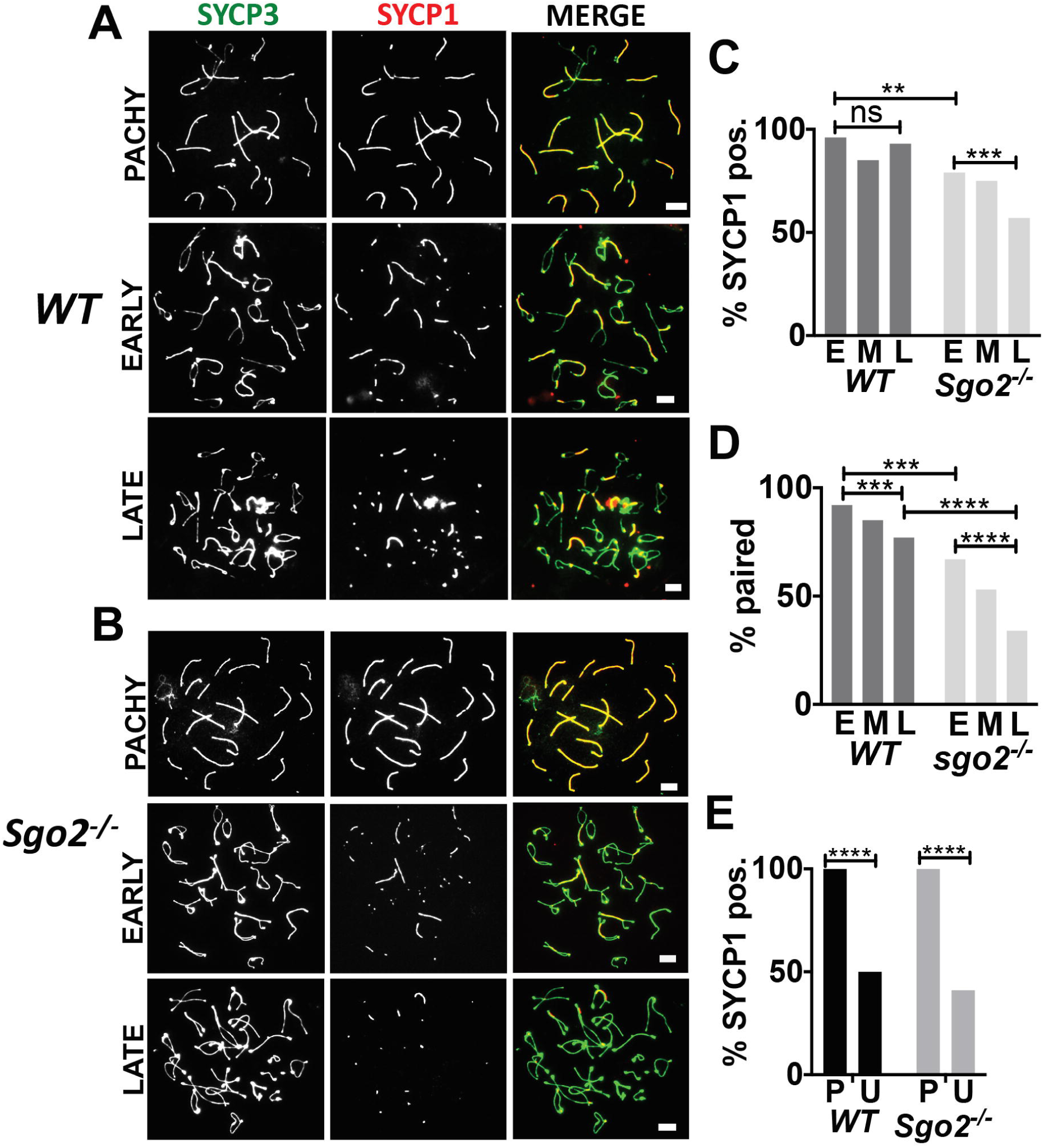
SGO2 is required for the persistence of centromeric synaptonemal complex components in diplotene. Indirect immunofluorescence was used to monitor the morphology of chromosomes from wild-type and *Sgo2* ^−/−^ spermatocytes. Representative chromosome spreads from pachytene, early diplotene and late diplotene (A) wild-type spermatocytes and (B) *Sgo2*^−/−^ spermatocytes. Scale bars represent 5 µm. (C) Histogram of SYCP1 localization at centromeres in spreads from wildtype and *Sgo2* ^−/−^ spermatocytes in early, E, middle, M, and late diplotene, L. The number of SYCP1 positive centromeres scored was: *WT* early (50/52), *WT* middle (46/54), *WT* late 88/95, *Sgo2* ^−/−^ (72/91) *Sgo2* ^−/−^ (114/151) *Sgo2* ^−/−^ (100/174). (D) Histogram of the percent paired centromeres on chromosomes from wildtype and *Sgo2* ^−/−^ spermatocytes. The number of paired centromeres was: *WT* early (48/52), *WT* middle (46/54), *WT* late (67/98), *Sgo2* ^−/−^ (61/91) *Sgo2* ^−/−^ (80/151) *Sgo2* ^−/−^ (59/174). (E) SYCP1 localization to paired and unpaired centromeres. (E) Paired (P) and unpaired (U) centromeres from all stages of diplotene (D above) were classified as according to their SYCP1 staining. The numbers of centromeres scored was*: WT* paired (167/167), *WT* unpaired (17/34*), Sgo2* ^−/−^ paired (200/200), *Sgo2* ^−/−^ unpaired (89/216). The significance of differences between samples was evaluated using Fisher’s exact test. **p < 0.01, *** p < 0.001, ****p < 0.0001.

The heightened loss of SYCP1 from centromeres would predict that *Sgo2^−/−^* spermatocytes would also have a defect in homologous centromere pairing in diplotene. In early diplotene, nearly all centromeres are paired in wild-type spermatocytes and pairing levels go down as cells proceed through diplotene (Fig. 3 A and D). In contrast, by early diplotene in *Sgo2*^−/−^ cells many of the centromere pairs have already disengaged and pair levels go down through diplotene (Fig. 3 B, and D). In both the wild-type control and the *Sgo2*^−/−^ chromosome spreads, the unpaired centromeres have significantly less SYCP1 staining than do the paired centromeres (Fig. 3 E, Supl. Fig. 3), supporting the notion that it is the protection of SYCP1 that allows centromere pairing to persist.

Together these results suggest that wild-type and *Sgo2*^−/−^ spermatocytes have similar SC structures and complete centromere pairing in pachytene. Importantly, in early diplotene in *Sgo2*^−/−^ mutants, SC is present at most centromeres and they remain paired. Thus, SGO2 is not necessary for establishing centromere pairing. However, centromere pairing disappears more rapidly in *Sgo2^−/−^* spermatocytes suggesting that SGO2 is necessary to maintain centromeric SC components, and centromere pairing, in diplotene.

### PP2A promotes centromere pairing in mouse spermatocytes

SGO2 could be protecting centromeric SC through the recruitment of one of its effector proteins to the centromere (reviewed in (22)). Shugoshin recruits PP2A phosphatase to meiotic centromeres in germ cells, where the PP2A opposes the phosphorylation of centromeric cohesins (23, 24, 47). To test whether this mechanism is being used to protect centromeric SC from disassembly in diplotene, we evaluated the persistence of centromeric SC, and centromere pairing, in spermatocytes when phosphatase activity was inhibited (Fig. 4). In these experiments we evaluated diplotene-like chromosome spreads from cultured spermatocytes (48) treated with the phosphatase inhibitors cantharidin and okadaic acid, which at the concentrations used preferentially inhibit PP2A over other phosphatases (49, 50). Treatment with either inhibitor significantly reduced the retention of SYCP1 at the centromeres and resulted in a substantial loss of centromere pairing (Fig. 4 A-D), consistent with the model that PP2A protects SYCP1 at centromeres, although it is formally possible that other targets of these compounds could be involved.

**Figure 4.**
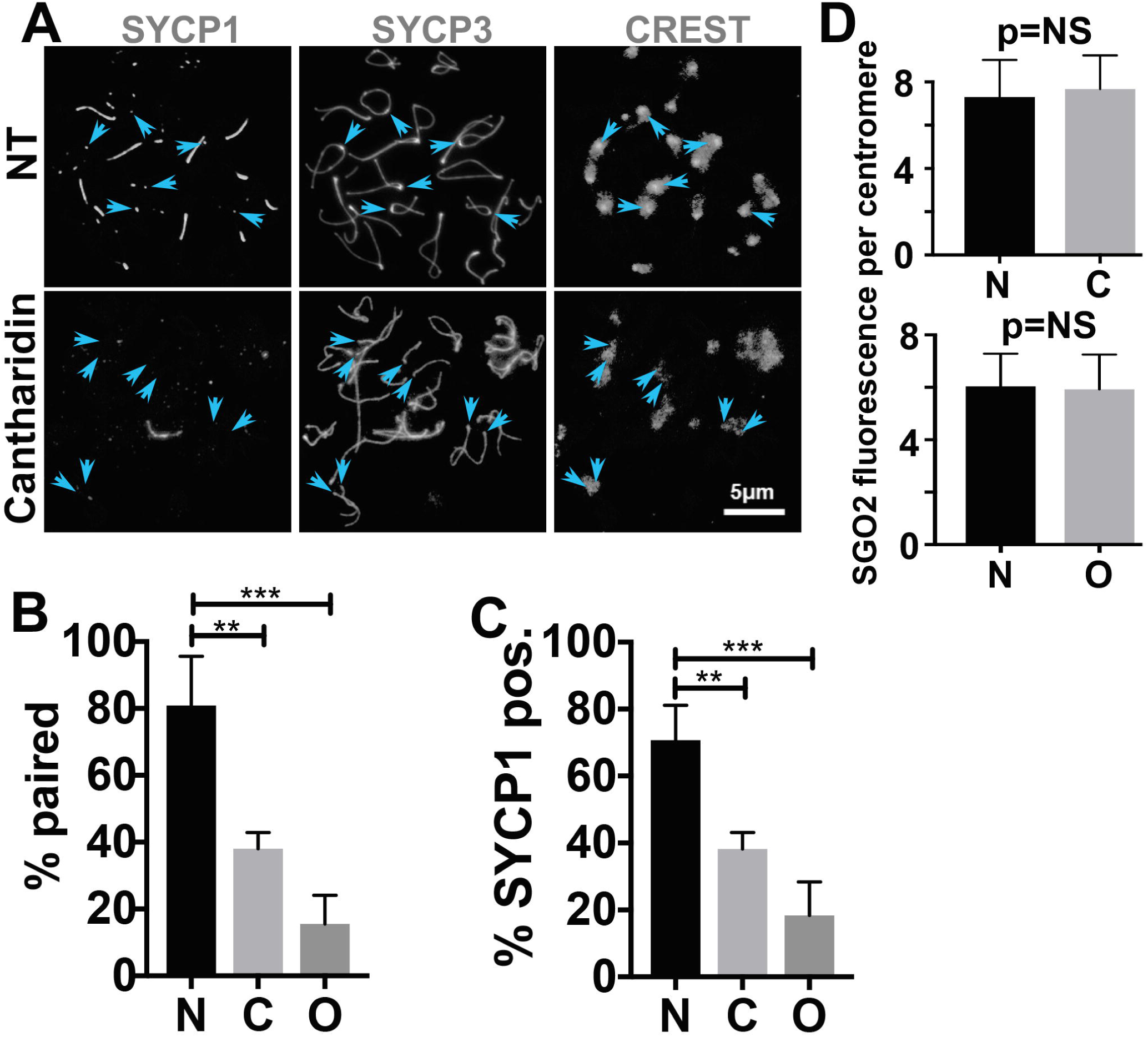
Phosphatase activity is needed for SYCP1 to persist at centromeres in diplotene. Cultured spermatocytes were treated with the phosphatase inhibitors okadaic acid or cantharidin. Chromosome spreads from diplotene cells were then evaluated using indirect immunofluorescence microscopy. The presence of SYCP1 at centromeres and the fraction of chromosomes in each spread with paired centromeres (stained with CREST antibodies) were scored. (A) Representative images of chromosome spreads that were not treated (NT) or treated with cantharidin. Scale bar represents 5 µm and applies to all panels. Arrowheads indicate the locations of examples of paired centromeres in top panels and unpaired centromeres in bottom panel. One hundred diplotene chromosome spreads were scored for SCYP1 localization to centromeres and centromere pairing. (B) The percentage of chromosomes in each spread with SYCP1 at the centromeres. Averages and standard deviations are: N (not treated) 80.9 +/−14.7%. C (cantharidin) 38.0+/−4.9%. O (okadaic acid) 15.5+/8.6%. (C) The percentage of chromosomes in each spread with paired centromeres. Averages and standard deviations are: N (not treated) 70.7 +/−10.4%. C (cantharidin) 38.2+/− 4.9%. O (okadaic acid) 18.4+/10.0%. One hundred chromosome spreads were scored for each treatment. Significance was evaluated using the student’s t test. (D). Histogram showing the relative amount of SGO2 on centromeres of untreated (N) or cantharidin (C) or okadaic acid (O) treated spermatocytes. ** p < 0.01, *** p < 0.001.

Recent studies in *Drosophila* have suggested that PP2A and shugoshin might each act to promote localization of the other to the centromeres (32), but in our experiments no reduction in SGO2 localization was seen at the centromeres following addition of the phosphatase inhibitors (Fig. 4 A and D). Although it is possible that both inhibitors are achieving their effects through some other target, the fact that both of these PP2A inhibitors reduce SC protection at centromeres is consistent with the model that PP2A, recruited by SGO2, opposes kinase activities that promote SC disassembly at centromeres upon pachytene exit.

### Shugoshin promotes disjunction of non-exchange chromosomes

In budding yeast, it has been possible to demonstrate directly that centromere pairing in prophase is necessary for subsequent disjunction of non-exchange partner chromosomes in anaphase I (51). Since shugoshin is acting at the centromeres in this interval we tested whether it is important in promoting the meiotic segregation of non-exchange partner chromosomes. In the mouse model, there is no established system for following the fate of non-exchange partner chromosomes, so we addressed this question using budding yeast, which has a single shugoshin gene, *SGO1*. Yeast *sgo1Δ* (deletion) mutants show only low levels of meiosis I non-disjunction (of exchange chromosomes), but severe defects in meiosis II (52). As was first shown in *Drosophila* (20, 21), the meiosis II defect is due to a failure to protect centromeric cohesion at the metaphase I to anaphase I transition. We first monitored the requirement for *SGO1* for centromere pairing using a pair of centromere-containing plasmids that act as non-exchange mini-chromosome partners in meiosis (53). These mini-chromosomes do not experience exchanges, yet they disjoin properly in most meioses (4, 54, 55). Cells bearing the mini-chromosomes were induced to enter meiosis, chromosome spreads were prepared, and pachytene spreads (as judged by Zip1 morphology) were scored for the association of the two mini-chromosomes, which were tagged at their centromeres with GFP and tdTomato (Fig. 5 A). The distances between the red and green foci marking the mini-chromosomes was measured in wild-type (*SGO1*) cells and cells that do not express *SGO1* in meiosis (*sgo1-md*) (56). Foci with center-to-center distances of less than 0.6 μm were as scored as “paired” (as in Fig. 5 A top panel) while those farther apart were scored as “unpaired” (as in Fig. 5 A, bottom panel). The *sgo1-md* mutants show considerable centromere pairing in pachytene, though at slightly lower levels than the control (Fig. 5 B). Deletion of the SC gene *ZIP1*, whose protein is thought to mediate centromere pairing, reduces pairing to a few percent in these assays (53). Thus, as was seen in mice (Fig. 3) shugoshin is not essential for establishing centromere paring. To test whether Sgo1 is necessary for the persistence of centromere pairing after pachytene exit, as it is in spermatocytes (Fig. 3 D), cells were synchronously released from a pachytene arrest and centromere pairing was scored at timed intervals. The arrest/release was achieved by using strains in which the *NDT80* meiotic transcription factor was under the control of an estradiol-inducible promotor (36, 57). Following pachytene release cells synchronously pass through diplotene and by two hours begin entering early metaphase (Supl. Fig, 4). The centromere paring of mini-chromosomes tagged with mTurquoise or mVenus was scored using fluorescence microscopy (Fig. 5 C and D). In pachytene cells (just prior to addition of estradiol) centromeres were paired in most cells in both wild-type and *sgo1* mutants (Fig. 5 D). As in the spermatocytes, centromere pairing levels diminished as cells continued meiotic progression (Fig. 5 D). By early metaphase the *sgo1* mutants had significantly lower levels of pairing than the wild-type cells (Fig. 5 D). Therefore, as in mouse spermatocytes, centromere pairing is naturally lost following pachytene and shugoshin slows the loss of pairing after pachytene exit.

**Figure 5.**
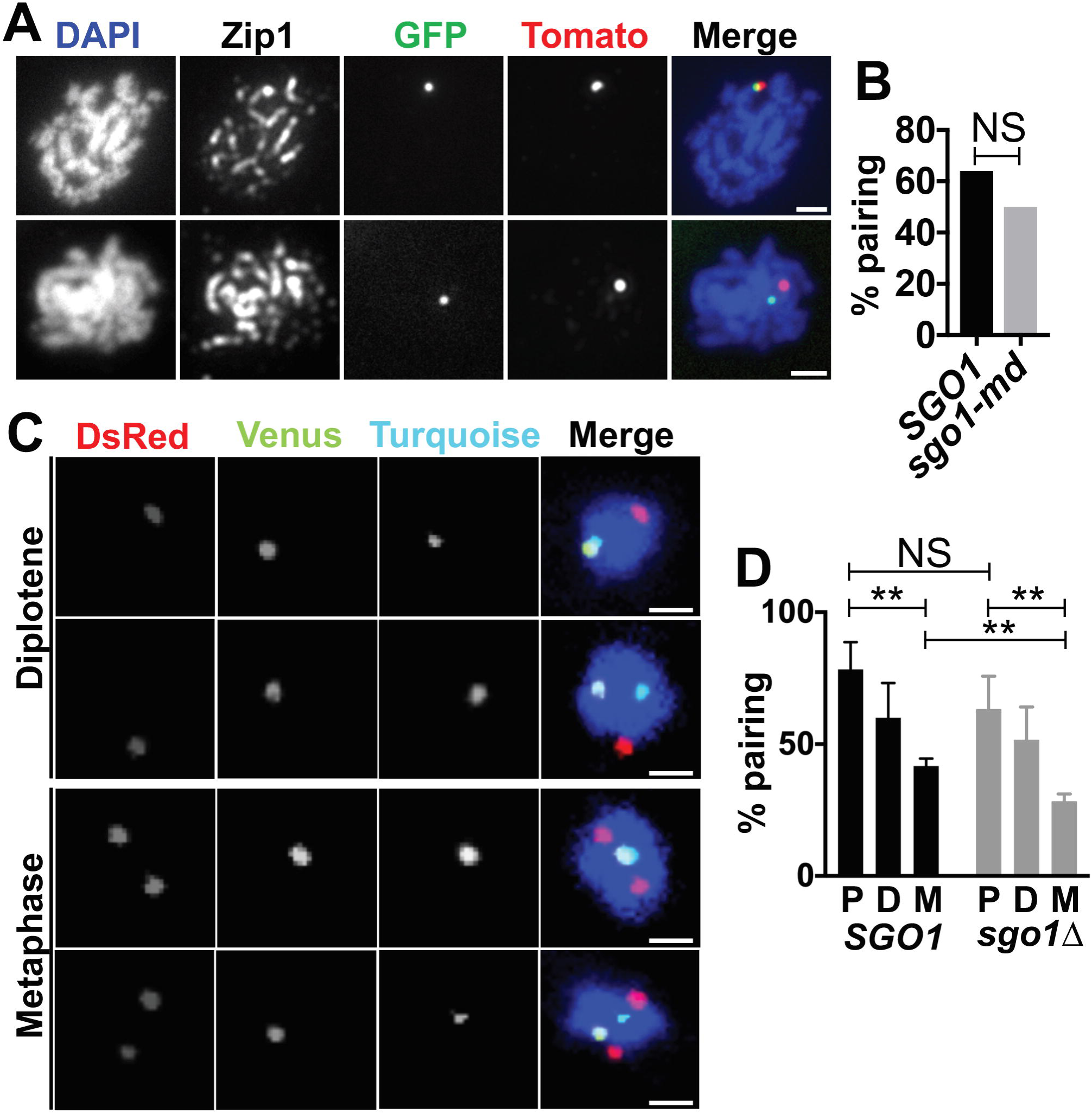
Shugoshin is not needed for centromere pairing in pachytene but is required for non-exchange segregation. (A) Representative chromosome spreads showing examples of paired (top) and unpaired (bottom) mini-chromosome centromeres (both images from the *sgo1-md* strain). Chromosome spreads were stained with DAPI to show chromatin, anti-Zip1 antibody to show the SC, and anti-GFP and DsRed to show the locations of the centromere-proximal tags. Scale bar equals 1 μm. (B) Histogram showing percent centromere pairing in each strain (*SGO1* n = 50 spreads, *sgo1-md* n = 100 spreads). (C) Representative micrographs showing examples of paired (top) and unpaired (bottom) mini-chromosome centromeres for both diplotene and metaphase cells (all images from the *SGO1* strain DJE90). Cells expressed mVenus-lacI and tetR-mTurquoise which bound to *lac* operator and *tet* operator arrays that were adjacent to the centromeres on the two mini-chromosomes. Spindle pole bodies are shown in red (*SPC42-DSRed*). Chromosomes were stained with DAPI. Scale bar equals 1μm. (D) Histogram showing percent pairing in pachytene, diplotene, and early-metaphase cells (P,D, and M, respectively) in *SGO1* and *sgo1Δ* cells. Experiments were performed in three replicates of 20 cells each. Statistical comparisons were performed with an unpaired t-test. For all histograms, NS = not significant and ** p < 0.01.

In both spermatocytes and budding yeast, centromere pairing occurs in prophase and in yeast it is necessary for disjunction in anaphase (14, 15) – even though the pairing has largely dissolved well before anaphase (Fig. 5 D). We have shown the centromere pairing is largely intact in shugoshin mutants but shugoshin is affecting centromere biology after pachytene exit, as centromere pairing and SC structures at centromeres are lost faster in shugoshin mutants.

To determine whether shugoshin is involved in later events in non-exchange centromere behavior, we examined the segregation of non-exchange mini-chromosomes in anaphase I (Fig. 6 A). Fluorescence microscopy was used to determine whether mini-chromosomes (marked by tdTomato and GFP foci) segregated to opposite poles in anaphase I cells. In the wild-type control, the mini-chromosomes non-disjoined in about 26% of meioses while in the *sgo1-md* mutants they non-disjoined in approximately 50% of meioses – consistent with random segregation (Fig. 6 B). Thus, although most mini-chromosomes pair at their centromeres in pachytene in *sgo1* mutants (Fig. 5 D), the pairing does not ensure disjunction. We also tested the role of Sgo1 in promoting the disjunction of authentic yeast chromosomes. In these experiments the yeast carried either a normal chromosome *V* pair or a pair of homeologous chromosomes *V’*s (one from *S. cerevisiae* and one from *S. bayanus*) that do not experience crossovers in meiosis because of sequence divergence (58). Both chromosome pairs were tagged with GFP at the centromeres. Deletion of *SGO1* (*sgo1Δ*) resulted in a small increase in non-disjunction frequency of the homologous chromosome *V*s (Fig. 6 C), consistent with earlier studies (52). In contrast, the non-exchange, homeologous, pair exhibited a significant increase in non-disjunction when *SGO1* was deleted, consistent with nearly random segregation (Fig. 6 D).

**Figure 6.**
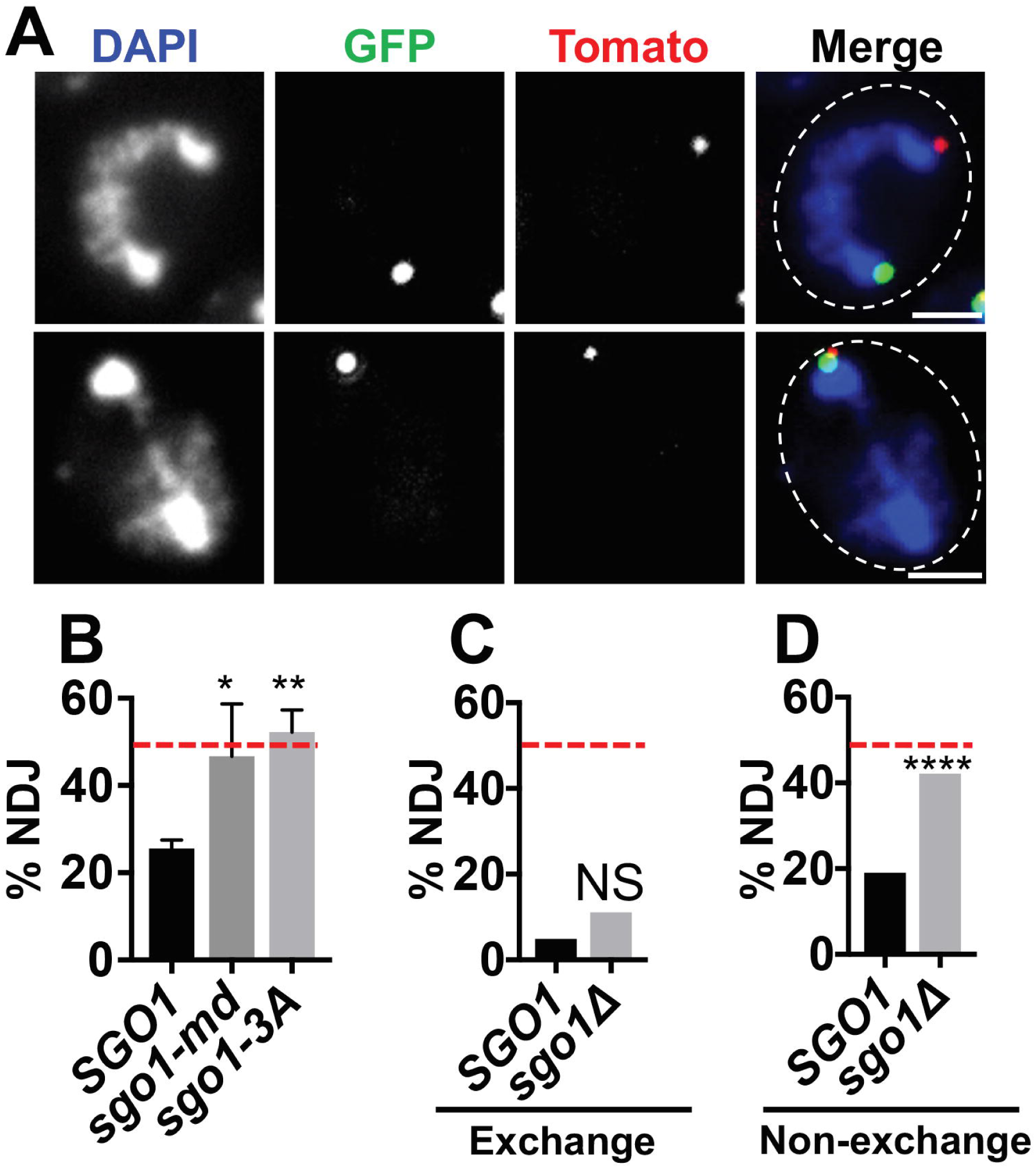
Shugoshin is required for non-exchange segregation of mini-chromosomes. (A) Representative anaphase cells showing disjoined (top) and non-disjoined (bottom) mini-chromosomes. Cells were stained with DAPI to show chromatin. Locations of mini-chromosome centromeres were detected by GFP and tdTomato fluorescence. Scale bar equals 1 μm. (B) Histogram showing non-disjunction frequencies of mini-chromosomes in *SGO1*, *sgo1-md*, and *sgo1-3A* cells (three replicates of 30 cells each were analyzed for all three strains). Statistical comparisons were performed using an unpaired t-test. (C) Histogram showing non-disjunction frequencies of homologous chromosomes in *SGO1* and *sgo1Δ* cells. (n = 122 cells for *SGO1* and 90 for *sgo1Δ*). (D) Histogram showing non-disjunction frequencies of homeologous Δ chromosomes in *SGO1* and *sgo1Δ* cells. (n = 121 cells for both strains, non-disjunction frequencies were 19.0% vs 42.1%, p < 0.0001). (B, C, D) Red line equals the level of non-disjunction expected for random segregation. (C,D) Statistical comparisons were performed with Fisher’s exact test. For all histograms, NS = not significant, * p < 0.05, ** p < 0.01, **** p < 0.0001.

The PP2A inhibitor experiments suggested that Sgo2 (in mice) acts to protect centromere pairing through recruitment of PP2A. Is Sgo1 of yeast promoting non-exchange disjunction by recruiting PP2A? To test this, we took advantage of the *sgo1-3A* allele in which three critical contact amino acids required for Sgo1 to recruit PP2A to centromeres are converted to alanines (27). This mutant, which exhibits normal loading of Sgo1 to meiotic kinetochores (27), showed random segregation of the non-exchange partners, providing strong support for the model that Sgo1 promotes non-exchange segregation through the recruitment of PP2A to centromeres.

These experiments show that shugoshin is not needed for the establishment of centromere pairing, that centromere pairing dissolves after pachytene exit, and that shugoshin slows the dissolution of centromere pairing. While previous work demonstrated that prophase centromere pairing is essential for non-exchange disjunction in yeast (14, 15), these results reveal that centromere pairing is not sufficient to ensure disjunction – since in yeast *SGO1* mutants most centromeres are paired upon pachytene exit. Earlier studies found that in wild-type cells, centromeric SC proteins disappear before chromosomes begin to orient on the spindle in early metaphase (14–17). This observation, coupled with our findings, suggests that centromere pairing might be a pre-condition for setting up the mechanism that later promotes bi-orientation in anaphase. Centromere pairing cannot be directly mediating bi-orientation since SC proteins are normally gone from the centromeres, and centromere paring is dissolved, before bi-orientation is accomplished. The observation that non-exchange partners in *Drosophila* appear to be tethered by threads of peri-centromeric heterochromatin during the metaphase bi-orientation process (59) suggests the model that prophase centromere pairing could provide a platform for the establishment of centromeric connections between non-exchange partners. The fact that non-exchange segregation is randomized in yeast shugoshin mutants raises the possibility that shugoshin is not essential for centromere pairing, *per se*, but instead the formation or maintenance of a structure or process that promotes bi-orientation. By this model, the low levels of meiosis I non-disjunction of native chromosomes in shugoshin mutants may reflect the times at which these chromosomes fail to experience exchanges and rely upon a centromere pairing based mechanism to ensure their disjunction in meiosis I.

## MATERIALS AND METHODS

The mouse strains, yeast strains, reagents, and methods for the experiments in this project are described in detail in the Supporting Information Appendix. The Oklahoma Medical Research Foundation Animal Care and Use Committee approved all animal protocols. For mouse cytology experiments, staging of chromosome spreads in diplotene was based on the extent of SYCP1 staining. Established approaches were employed for visualizing chromosomes in surface spreads in mice and yeast (16, 43, 60, 61).

## Acknowledgements

The authors wish to thank the members of the Program in Cell Cycle and Cancer Biology for their constructive comments during the course of this project. AMP is funded by BFU2017-89408-R from the Spanish Ministry of Economy (MINECO). RJP was supported by COBRE grant GM103636, and March of Dimes grant FY14-256. DSD was supported by NIH grant R01 GM087377.

## References

1. Hunter N (2015) Meiotic Recombination: The Essence of Heredity. Cold Spring Harb Perspect Biol 7(12).

2. Kurdzo EL & Dawson DS (2015) Centromere pairing--tethering partner chromosomes in meiosis I. FEBS J 282(13):2458–2470.

3. Carpenter AT (1973) A meiotic mutant defective in distributive disjunction in Drosophila melanogaster. Genetics 73(3):393–428.

4. Dawson DS, Murray AW, & Szostak JW (1986) An alternative pathway for meiotic chromosome segregation in yeast. Science 234(4777):713–717.

5. Guacci V & Kaback DB (1991) Distributive disjunction of authentic chromosomes in Saccharomyces cerevisiae. Genetics 127(3):475–488.

6. Hawley RS, et al. (1992) There are two mechanisms of achiasmate segregation in Drosophila females, one of which requires heterochromatic homology. Dev Genet 13(6):440–467.

7. Woods LM, et al. (1999) Chromosomal influence on meiotic spindle assembly: abnormal meiosis I in female Mlh1 mutant mice. The Journal of cell biology 145(7):1395–1406.

8. Cheng EY, et al. (2009) Meiotic recombination in human oocytes. PLoS Genet 5(9):e1000661.

9. Oliver TR, et al. (2008) New insights into human nondisjunction of chromosome 21 in oocytes. PLoS Genet 4(3):e1000033.

10. Fledel-Alon A, et al. (2009) Broad-scale recombination patterns underlying proper disjunction in humans. PLoS Genet 5(9):e1000658.

11. Tease C, Hartshorne GM, & Hulten MA (2002) Patterns of meiotic recombination in human fetal oocytes. American journal of human genetics 70(6):1469–1479.

12. Dernburg AF, Sedat JW, & Hawley RS (1996) Direct evidence of a role for heterochromatin in meiotic chromosome segregation. Cell 86(1):135–146.

13. Ding DQ, Yamamoto A, Haraguchi T, & Hiraoka Y (2004) Dynamics of homologous chromosome pairing during meiotic prophase in fission yeast. Dev Cell 6(3):329–341.

14. Gladstone MN, Obeso D, Chuong H, & Dawson DS (2009) The synaptonemal complex protein Zip1 promotes bi-orientation of centromeres at meiosis I. PLoS Genet 5(12):e1000771.

15. Newnham L, Jordan P, Rockmill B, Roeder GS, & Hoffmann E (2010) The synaptonemal complex protein, Zip1, promotes the segregation of nonexchange chromosomes at meiosis I. Proc Natl Acad Sci U S A 107(2):781–785.

16. Bisig CG, et al. (2012) Synaptonemal complex components persist at centromeres and are required for homologous centromere pairing in mouse spermatocytes. PLoS Genet 8(6):e1002701.

17. Qiao H, et al. (2012) Interplay between synaptonemal complex, homologous recombination, and centromeres during mammalian meiosis. PLoS Genet 8(6):e1002790.

18. Takeo S, Lake CM, Morais-de-Sa E, Sunkel CE, & Hawley RS (2011) Synaptonemal complex-dependent centromeric clustering and the initiation of synapsis in Drosophila oocytes. Curr Biol 21(21):1845–1851.

19. Rankin S (2015) Complex elaboration: making sense of meiotic cohesin dynamics. FEBS J 282(13):2426–2443.

20. Kerrebrock AW, Miyazaki WY, Birnby D, & Orr-Weaver TL (1992) The Drosophila mei-S332 gene promotes sister-chromatid cohesion in meiosis following kinetochore differentiation. Genetics 130(4):827–841.

21. Kerrebrock AW, Moore DP, Wu JS, & Orr-Weaver TL (1995) Mei-S332, a Drosophila protein required for sister-chromatid cohesion, can localize to meiotic centromere regions. Cell 83(2):247–256.

22. Marston AL (2015) Shugoshins: tension-sensitive pericentromeric adaptors safeguarding chromosome segregation. Mol Cell Biol 35(4):634–648.

23. Kitajima TS, et al. (2006) Shugoshin collaborates with protein phosphatase 2A to protect cohesin. Nature 441(7089):46–52.

24. Lee J, et al. (2008) Unified mode of centromeric protection by shugoshin in mammalian oocytes and somatic cells. Nat Cell Biol 10(1):42–52.

25. Rattani A, et al. (2013) Sgol2 provides a regulatory platform that coordinates essential cell cycle processes during meiosis I in oocytes. Elife 2:e01133.

26. Riedel CG, et al. (2006) Protein phosphatase 2A protects centromeric sister chromatid cohesion during meiosis I. Nature 441(7089):53–61.

27. Xu Z, et al. (2009) Structure and function of the PP2A-shugoshin interaction. Mol Cell 35(4):426–441.

28. Ishiguro T, Tanaka K, Sakuno T, & Watanabe Y (2010) Shugoshin-PP2A counteracts casein-kinase-1-dependent cleavage of Rec8 by separase. Nat Cell Biol 12(5):500–506.

29. Katis VL, et al. (2010) Rec8 phosphorylation by casein kinase 1 and Cdc7-Dbf4 kinase regulates cohesin cleavage by separase during meiosis. Dev Cell 18(3):397–409.

30. Rumpf C, et al. (2010) Casein kinase 1 is required for efficient removal of Rec8 during meiosis I. Cell Cycle 9(13):2657–2662.

31. Cahoon CK & Hawley RS (2016) Regulating the construction and demolition of the synaptonemal complex. Nat Struct Mol Biol 23(5):369–377.

32. Jordan PW, Karppinen J, & Handel MA (2012) Polo-like kinase is required for synaptonemal complex disassembly and phosphorylation in mouse spermatocytes. J Cell Sci 125(Pt 21):5061–5072.

33. Tarsounas M, Pearlman RE, & Moens PB (1999) Meiotic activation of rat pachytene spermatocytes with okadaic acid: the behaviour of synaptonemal complex components SYN1/SCP1 and COR1/SCP3. J Cell Sci 112 (Pt 4):423–434.

34. Sourirajan A & Lichten M (2008) Polo-like kinase Cdc5 drives exit from pachytene during budding yeast meiosis. Genes Dev 22(19):2627–2632.

35. Argunhan B, et al. (2017) Fundamental cell cycle kinases collaborate to ensure timely destruction of the synaptonemal complex during meiosis. EMBO J 36(17):2488–2509.

36. Carlile TM & Amon A (2008) Meiosis I is established through division-specific translational control of a cyclin. Cell 133(2):280–291.

37. Matos J, et al. (2008) Dbf4-dependent CDC7 kinase links DNA replication to the segregation of homologous chromosomes in meiosis I. Cell 135(4):662–678.

38. Sun F & Handel MA (2008) Regulation of the meiotic prophase I to metaphase I transition in mouse spermatocytes. Chromosoma 117(5):471–485.

39. Anderson LK, Reeves A, Webb LM, & Ashley T (1999) Distribution of crossing over on mouse synaptonemal complexes using immunofluorescent localization of MLH1 protein. Genetics 151(4):1569–1579.

40. Jagiello G & Fang JS (1979) Analyses of diplotene chiasma frequencies in mouse oocytes and spermatocytes in relation to ageing and sexual dimorphism. Cytogenet Cell Genet 23(1-2):53–60.

41. Speed RM (1977) The effects of ageing on the meiotic chromosomes of male and female mice. Chromosoma 64(3):241–254.

42. Zelazowski MJ, et al. (2017) Age-Dependent Alterations in Meiotic Recombination Cause Chromosome Segregation Errors in Spermatocytes. Cell 171(3):601–614 e613.

43. Guiraldelli MF, Eyster C, Wilkerson JL, Dresser ME, & Pezza RJ (2013) Mouse HFM1/Mer3 is required for crossover formation and complete synapsis of homologous chromosomes during meiosis. PLoS Genet 9(3):e1003383.

44. Lipkin SM, et al. (2002) Meiotic arrest and aneuploidy in MLH3-deficient mice. Nat Genet 31(4):385–390.

45. Holloway JK, Booth J, Edelmann W, McGowan CH, & Cohen PE (2008) MUS81 generates a subset of MLH1-MLH3-independent crossovers in mammalian meiosis. PLoS Genet 4(9):e1000186.

46. Gomez R, et al. (2007) Mammalian SGO2 appears at the inner centromere domain and redistributes depending on tension across centromeres during meiosis II and mitosis. EMBO Rep 8(2):173–180.

47. Llano E, et al. (2008) Shugoshin-2 is essential for the completion of meiosis but not for mitotic cell division in mice. Genes Dev 22(17):2400–2413.

48. La Salle S, Sun F, & Handel MA (2009) Isolation and short-term culture of mouse spermatocytes for analysis of meiosis. Methods Mol Biol 558:279–297.

49. Honkanen RE (1993) Cantharidin, another natural toxin that inhibits the activity of serine/threonine protein phosphatases types 1 and 2A. FEBS Lett 330(3):283–286.

50. Bialojan C & Takai A (1988) Inhibitory effect of a marine-sponge toxin, okadaic acid, on protein phosphatases. Specificity and kinetics. Biochem J 256(1):283–290.

51. Kurdzo EL, Chuong HH, Evatt JM, & Dawson DS (2018) A ZIP1 separation-of-function allele reveals that centromere pairing drives meiotic segregation of achiasmate chromosomes in budding yeast. PLoS Genet 14(8):e1007513.

52. Marston AL, Tham WH, Shah H, & Amon A (2004) A genome-wide screen identifies genes required for centromeric cohesion. Science 303(5662):1367–1370.

53. Kurdzo EL, Obeso D, Chuong H, & Dawson DS (2017) Meiotic Centromere Coupling and Pairing Function by Two Separate Mechanisms in Saccharomyces cerevisiae. Genetics 205(2):657–671.

54. Mann C & Davis RW (1986) Meiotic disjunction of circular minichromosomes in yeast does not require DNA homology. Proc Natl Acad Sci U S A 83(16):6017–6019.

55. Sears DD, Hegemann JH, & Hieter P (1992) Meiotic recombination and segregation of human-derived artificial chromosomes in Saccharomyces cerevisiae. Proc Natl Acad Sci U S A 89(12):5296–5300.

56. Monje-Casas F, Prabhu VR, Lee BH, Boselli M, & Amon A (2007) Kinetochore orientation during meiosis is controlled by Aurora B and the monopolin complex. Cell 128(3):477–490.

57. Benjamin KR, Zhang C, Shokat KM, & Herskowitz I (2003) Control of landmark events in meiosis by the CDK Cdc28 and the meiosis-specific kinase Ime2. Genes Dev 17(12):1524–1539.

58. Maxfield Boumil R, Kemp B, Angelichio M, Nilsson-Tillgren T, & Dawson DS (2003) Meiotic segregation of a homeologous chromosome pair. Mol Genet Genomics 268(6):750–760.

59. Hughes SE, et al. (2009) Heterochromatic threads connect oscillating chromosomes during prometaphase I in Drosophila oocytes. PLoS Genet 5(1):e1000348.

60. Kemp B, Boumil RM, Stewart MN, & Dawson DS (2004) A role for centromere pairing in meiotic chromosome segregation. Genes Dev 18(16):1946–1951.

61. Handel MA, Caldwell KA, & Wiltshire T (1995) Culture of pachytene spermatocytes for analysis of meiosis. Dev Genet 16(2):128–139.

