## Supporting Information for "Shugoshin protects centromere pairing and promotes segregation of non-exchange partner chromosomes in meiosis"

### MATERIALS AND METHODS

#### Yeast Strains

Yeast culturing and sporulation was as described previously (1). Strains are isogenic derivatives of rapidly sporulating strains of primarily SK1 and W303 ancestry, derived in the RE Esposito laboratory (2). Complete sequences of the parental strains are available at NCBI.

#### Mouse Strains

The Oklahoma Medical Research Foundation Animal Care and Use Committee approved all animal protocols. The following mice were used in this study: C57BL/6, Shugoshin $2^{-/-}$ (3), $Hfm1^{-/-}$ (4), and $Mus81^{-/-}$ (5), a gift from Paula Cohen.

#### Spermatocyte Cytology

Established approaches were employed for visualizing chromosomes in surface spreads (4, 6). Fixed spermatocyte images were analyzed using AxioVision software (Zeiss). Statistical tests were as described in the figure legends or text. For staging spermatocyte chromosome spreads, those with continuous SYCP1 signal and completely synapsed SYCP3 staining lateral elements were scored as pachytene. Staging of chromosome spreads in diplotene was based on the extent of SYCP1 staining. Those with five or more stretches of SYCP1 5 μm or longer were classified as early diplotene, those with two-to-four such SYCP1 stretches were scored as middle diplotene, and those with one or fewer 5 μm runs of SYCP1 were classified as late diplotene. The percent of centromere pairing (Fig. 1, 3 and 4) was tabulated for chromosome ends that could clearly be resolved in the chromosome spread. The percent SYCP1 staining of centromeres was scored for pairs of centromeres (Fig.3 and 4). In this assay, for unpaired centromeres, if either of the centromeres exhibited overlapping SYCP1 staining the pair was scored as SYCP1 positive.

#### Spermatocyte culture and chemical inhibition

Short-term culture of spermatocytes was performed essentially as described (7-10). Okadaic acid and cantharidin were added at concentrations of 5 μM and 10 μM respectively. At these concentrations Okadaic acid and cantharidin are 10 – 100 times more potent inhibitors of PP2A than PP1 (11) (12). Equivalent volumes of DMSO or ethanol alone were added to “no treatment” control cultures. In all cell culture experiments, cell viability was quantified using the Trypan Blue exclusion assay.

#### Imaging

All images were collected using the 100X objective lens of a Zeiss AxioImager microscope with band-pass emission filters, a Roper HQ2 CCD. Image processing and measurements of image features were performed with, and AxioVision software.

#### Yeast Centromere Pairing In Pachytene Assay

Chromosome V centromere pairing in pachytene was evaluated using published methods (13) in which a *lac* operator array was inserted adjacent to the centromere of two chromosomes and a lacI-GFP hybrid protein was expressed under the control of a meiotic promoter to produce a focus of GFP at the *lac* operator arrays (14). Chromosome spreads were prepared and stained DAPI (4,6-Diamidino-2-phenylindole, dihydrochloride) to allow visualization of chromatin and with the primary antibodies mouse anti-Zip1p (a gift from Rebecca Maxfield), and chicken anti-GFP (Chemicon AB16901). Secondary antibodies were Alexa Fluor 488-conjugated goat anti-chicken IgG and Alexa Fluor 546-conjugated goat anti-mouse IgG (all from Molecular Probes). All were used at 1:1000 dilution. Immunofluorescence microscopy was used to identify those spreads with the condensed chromosomes typical of late meiotic prophase and the number and proximity of GFP and tdTomoto foci was used as a measure of pairing (13). Spreads with one focus, or two foci within 0.6 microns, were scored as paired. Those in which the foci were separated by a larger distance were scored as unpaired. Measurements were performed with AxioVision software. Pairing between centromere plasmids was performed similarly. Each plasmid carried an origin of replication (*ARS1*) and a 5.1 kb EcoRI fragment from chromosome *III* that includes the centromere. One plasmid was tagged with tdTomato-tetR hybrid proteins that localize to a *tet* operon operator array adjacent to the centromere (15), the other is tagged with GFP-lacI hybrid proteins that localize to a *lac* operon operator array adjacent to its centromere (14). Chromosome spreads were prepared as described in (16), with the following modifications: Cells were harvested 5-7 hours after induction of sporulation at 30°C. Primary antibodies were mouse anti-Zip1 (used at 1:1000 dilution), chicken anti-GFP (used at 1:500 dilution; Millipore AB16901), rabbit anti-DsRed (used at 1:1000-1:2000 dilution; Clontech 632496). Secondary antibodies were obtained from Thermo Fisher: Alexa Fluor 488-conjugated goat anti-chicken IgG (used at 1:1200 dilution), Alexa Fluor 568-conjugated goat anti-mouse IgG (1:1000), Alexa Fluor 647 conjugated goat anti-rabbit IgG (used at 1:1200 dilution). Only cells that exhibited “ropey” DAPI staining were scored in this assay, and were disqualified for assessment if there was more than one GFP focus or more than one tdTomato focus. In these cells, the distance between the center of the green focus and the center of the red focus was measured using AxioVision software.

**Yeast Centromere Pairing After Release From A Pachytene Arrest**

Cells bearing plasmids marked at their centromeres by mVenus-tetR or mTurquoise-lacI were arrested in pachytene using the $P_{GAL1}$-*NDT80*, *GAL4-ER* system (17, 18). Addition of estradiol to the medium was used to induce *NDT80* expression, which triggers pachytene exit. Samples were harvested at one-hour intervals beginning at estradiol addition (T=0). Cells were examined using fluorescent microscopy on a Zeiss Axioimager with a Roper Coolsnap CCD. Cells were then scored for the percent that exhibited centromere pairing (the mTurquoise and mVenus dots are touching). In cells that fail to enter meiosis the centromeres remain clustered at the spindle pole bodies. These cells were ignored in the analysis. Cells from the T=0 timepoint were used to assess centromere pairing in pachytene, those at the T=1 hour timepoint were used to assess centromere pairing in diplotene, those at T=2 hours with spindles shorter than 1.75 microns (new spindles) were used to score pairing in pro-metaphase.

**Achiasmate segregation assay**

Non-disjunction frequencies of centromere plasmids were determined using published assays (19). Harvested cells were either assayed fresh or were frozen in 15% glycerol and 1% potassium acetate until the time at which they were assayed. Preparation for assaying the cells included staining the cells with DAPI and then mounting the cells on agarose pads for viewing as described previously (20). Anaphase I cells were identified by the presence of two DAPI masses on either side of elongated cells, indicating that the chromosomes had segregated. To avoid scoring cells with duplicated or lost CEN plasmids, only cells with one GFP focus and one tdTomato focus were assayed. Segregation of the GFP-tagged chromosome V's was done using similar methods following a previously described protocol (13)

1. Kurdzo EL, Obeso D, Chuong H, & Dawson DS (2017) Meiotic Centromere Coupling and Pairing Function by Two Separate Mechanisms in Saccharomyces cerevisiae. *Genetics* 205(2):657-671.
2. Dresser ME, Ewing DJ, Harwell SN, Coody D, & Conrad MN (1994) Nonhomologous synapsis and reduced crossing over in a heterozygous paracentric inversion in Saccharomyces cerevisiae. *Genetics* 138(3):633-647.
3. Llano E*, et al.* (2008) Shugoshin-2 is essential for the completion of meiosis but not for mitotic cell division in mice. *Genes Dev* 22(17):2400-2413.
4. Guiraldelli MF, Eyster C, Wilkerson JL, Dresser ME, & Pezza RJ (2013) Mouse HFM1/Mer3 is required for crossover formation and complete synapsis of homologous chromosomes during meiosis. *PLoS Genet* 9(3):e1003383.
5. Holloway JK, Booth J, Edelmann W, McGowan CH, & Cohen PE (2008) MUS81 generates a subset of MLH1-MLH3-independent crossovers in mammalian meiosis. *PLoS Genet* 4(9):e1000186.
6. Bisig CG*, et al.* (2012) Synaptonemal complex components persist at centromeres and are required for homologous centromere pairing in mouse spermatocytes. *PLoS Genet* 8(6):e1002701.
7. Handel MA, Caldwell KA, & Wiltshire T (1995) Culture of pachytene spermatocytes for analysis of meiosis. *Dev Genet* 16(2):128-139.
8. La Salle S, Sun F, & Handel MA (2009) Isolation and short-term culture of mouse spermatocytes for analysis of meiosis. *Methods Mol Biol* 558:279-297.
9. Rao HB*, et al.* (2017) A SUMO-ubiquitin relay recruits proteasomes to chromosome axes to regulate meiotic recombination. *Science* 355(6323):403-407.
10. Wiltshire T, Park C, Caldwell KA, & Handel MA (1995) Induced premature G2/M-phase transition in pachytene spermatocytes includes events unique to meiosis. *Dev Biol* 169(2):557-567.
11. Honkanen RE (1993) Cantharidin, another natural toxin that inhibits the activity of serine/threonine protein phosphatases types 1 and 2A. *FEBS Lett* 330(3):283-286.
12. Bialojan C & Takai A (1988) Inhibitory effect of a marine-sponge toxin, okadaic acid, on protein phosphatases. Specificity and kinetics. *Biochem J* 256(1):283-290.
13. Kemp B, Boumil RM, Stewart MN, & Dawson DS (2004) A role for centromere pairing in meiotic chromosome segregation. *Genes Dev* 18(16):1946-1951.
14. Straight AF, Belmont AS, Robinett CC, & Murray AW (1996) GFP tagging of budding yeast chromosomes reveals that protein-protein interactions can mediate sister chromatid cohesion. *Curr Biol* 6(12):1599-1608.
15. Michaelis C, Ciosk R, & Nasmyth K (1997) Cohesins: chromosomal proteins that prevent premature separation of sister chromatids. *Cell* 91(1):35-45.

16. Grubb J, Brown MS, & Bishop DK (2015) Surface Spreading and Immunostaining of Yeast Chromosomes. *J Vis Exp* (102):e53081.
17. Benjamin KR, Zhang C, Shokat KM, & Herskowitz I (2003) Control of landmark events in meiosis by the CDK Cdc28 and the meiosis-specific kinase Ime2. *Genes Dev* 17(12):1524-1539.
18. Carlile TM & Amon A (2008) Meiosis I is established through division-specific translational control of a cyclin. *Cell* 133(2):280-291.
19. Gladstone MN, Obeso D, Chuong H, & Dawson DS (2009) The synaptonemal complex protein Zip1 promotes bi-orientation of centromeres at meiosis I. *PLoS Genet* 5(12):e1000771.
20. Kim S, Meyer R, Chuong H, & Dawson DS (2013) Dual mechanisms prevent premature chromosome segregation during meiosis. *Genes Dev* 27(19):2139-2146.

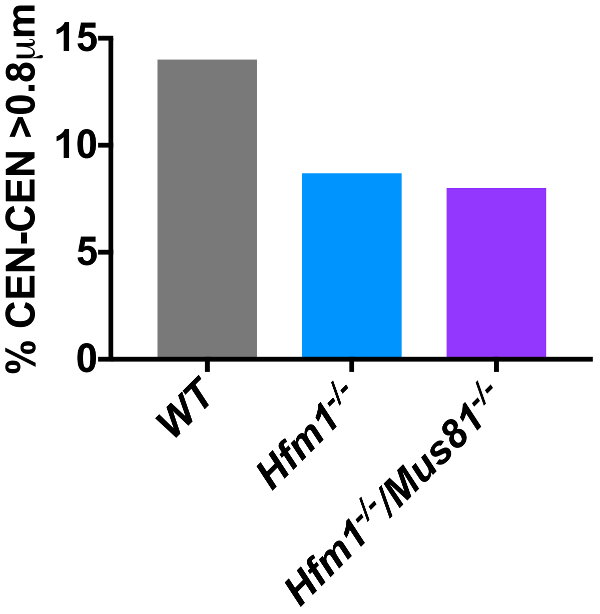


**Supplemental Figure 1. Centromere-centromere separation distances for achiasmate partners.** To test whether undetectable crossovers – those that do not produce a connection of the chromosome axes in a chromosome spread – we scored measured the distances between the centromeres of homologous partner chromosomes in diplotene chromosome spreads (Fig. 1 D). If there are undetectable crossovers that hold centromeres together in chromosome pairs with no apparent chiasma, then in recombination mutants, which should not have those crossovers, the centromeres of apparently achiasmate chromosomes should be free to separate. Here we re-plotted the data from Figure 1 D. Homologous centromeres farther apart than 0.8 microns were scored as “unpaired” as this is farther apart than the distance between the chromosome axes. The centromeres of homologous achiasmate chromosomes in the recombination mutants were no more likely to widely separated than in the WT cells, suggesting there are not undetectable crossovers connecting the centromeres of chromosomes that appear to be achiasmate in wildtype cells.

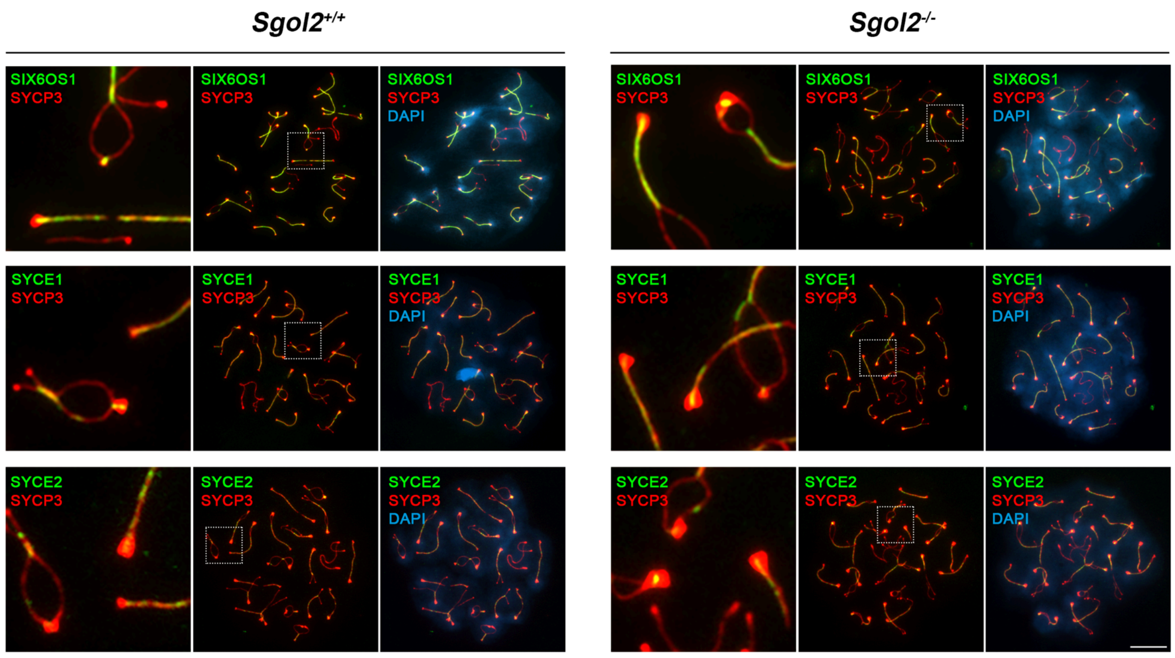


**Supplemental Figure 2. Synaptonemal complex proteins SIX60S1, SYCE1 and SYCE3 localize to paired centromeres.** Double immunolocalization of SYCP3 (red), and the central element components SIX6OS1 (upper panel), SYCE1 (central panel) and SYCE2 (lower panel) (green) in wild type and *Sgo2*$^{-/-}$ spermatocytes. DNA was stained with DAPI (blue). In the absence of SGO2, the three synaptonemal complex central element proteins persist at paired centromeres in early diplotene in a similar location as in wild type. Scale bars represent 10 μm.

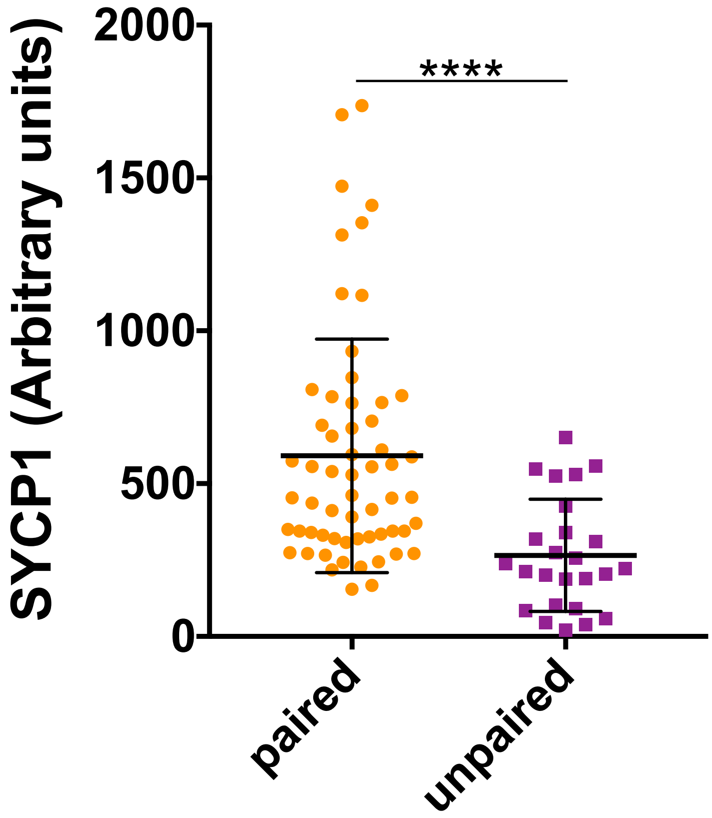


**Supplemental Figure 3. SYCP1 staining at paired and unpaired centromeres of achiasmate chromosomes.** The chromosomes spreads for wildtype cells in Figure 1 were analyzed to determine the relative staining intensities of SYCP1 at paired and unpaired centromeres. Axiovision software was used to mark an area encircling paired centromeres or two areas individually encircling unpaired centromeres using the CREST signal as a guide (see example in Fig. 1 A). The signal in the SYCP1 channel was recorded. The background signal from an equivalent area adjacent to the centromeres was obtained and subtracted from the SYCP1 value. The SYCP1 signals from the paired centromeres and the two summed values for each unpaired centromere pair are plotted. Paired n=59, unpaired n=25. Unpaired t test, p=0.0001. The amount of SYCP1 detected at unpaired centromeres was significantly less than the signal at paired centromeres, consistent with the hypothesis that reductions in SYCP1 levels at centromeres contribute to a loss of pairing.

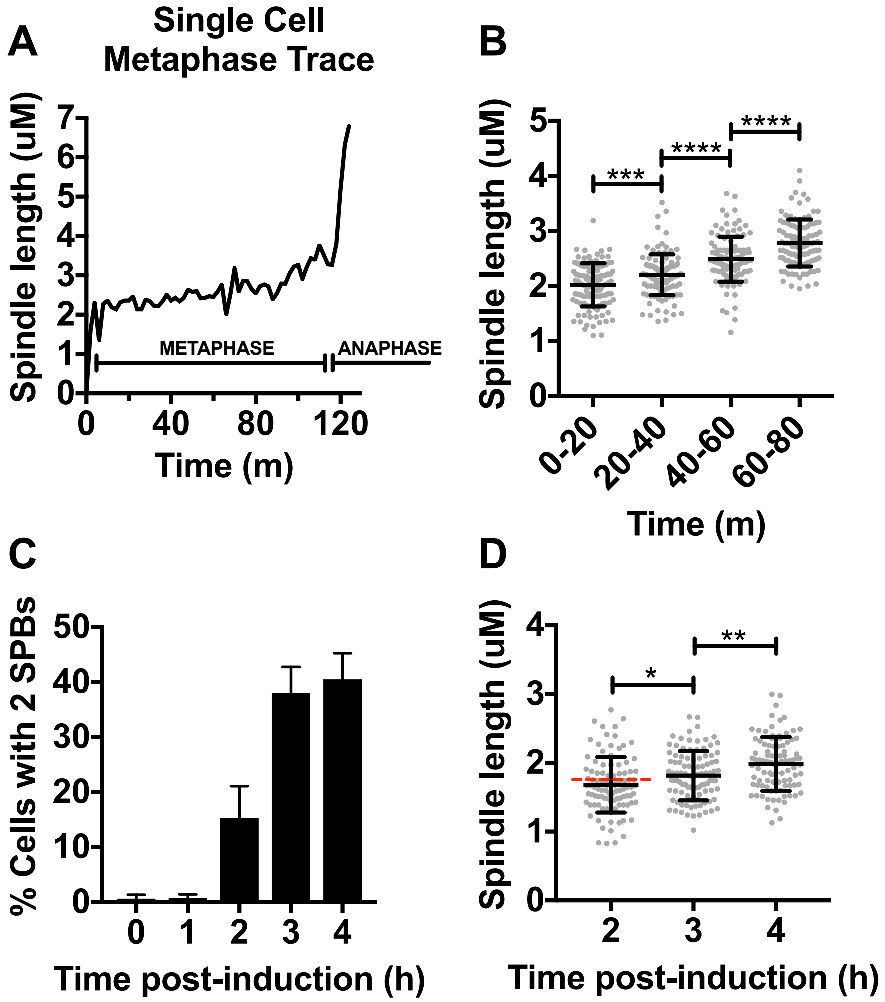


**Supplemental Figure 4. Spindle length increases during meiotic metaphase I. A.** Live cell imaging was used to track the distance between spindle poles on cells progressing through meiosis I. Spindle poles were marked with Spc42-DSRed (not shown). Imaging was performed on a Nikon Eclipse TE2000 at 63X acquiring images through five Z-planes every two minutes. Three-dimensional distances between the poles were measured using Nikon NIS Elements software. Shown is a trace of the distance between spindle poles starting shortly before the spindle poles separated (anaphase onset). **B.** Spindle length increases as metaphase progresses. The lengths of spindles were recorded every two minutes for each of twelve separate spindles. The pooled values for the twelve spindles, in bins of twenty minutes, were averaged. Length increases significantly over each twenty-minute interval in metaphase. **C.** Spindle formation after release from a pachytene arrest. Cells in the experiment shown in Figure 5 C-D were released from a pachytene arrest (see Materials and Methods). Cells were harvested at one-hour intervals. For each time point the percent of cells in which the spindle poles had separated to form a spindle were scored. Shown is an average of three experiments with n=200 cells per timepoint. New meiotic metaphase spindles were shown to begin forming starting at the two-hour timepoint. **D.** Spindle length increases with metaphase progression following release from pachytene arrest. The distance between spindle poles was measured in cells at two, three, and four hours after release from pachytene arrest. These measurements were only in the X-Y dimensions and are therefore a slight under-estimate of actual length as spindles might be tilted slightly in the Z dimension. Spindles increase in length significantly over time, consistent with newer spindles being shorter than older ones. Spindles shorter than 1.75uM, as indicated by the red dotted line, were classified as early metaphase spindles. For 2, 3, and 4 hours post-induction, n=99, 99, and 94 cells, respectively. Unpaired t-tests were used for statistical comparisons. *p<0.05, **p<0.01, ***p<0.001, ****p<0.0001.

**Table S1: Yeast Strains Used in This Study**

| Diploid Strains | Haploids | Name in Figure | Figure |
|---|---|---|---|
| DEK306 | X2187: *MAT***a**, *ura3-13, trp1-63, leu2, leu2::*OPL46[$P_{URA3}$*-tetR-tdTomato leu2::HIS3*], *met13-d, tyr1-1, can1-R, lys2::*pMDE798[$P_{DMC1}$*-GFP-lacI*], OPL210[*CEN3-5.1KB lacO256 LEU2 TRP1 ARS1*] OPL214[*CEN3-5.1KB, tetO256 URA3 TRP1 ARS1*] | *SGO1* | Fig. 5 B<br><br>Fig. 6 B |
| | Y2009: *MAT∝, trp1-Δ63, his3-Δ1, leu2, lys2::*pMDE798[$P_{DMC1}$*-GFP-lacI*], *tyr1-2, met13-c, cyh2-1* | | |
| DEK313 | X2351: *MAT***a**, *trp1-Δ63, his3-Δ1, leu2, met13-d, tyr1-1, lys2::*pMDE798[$P_{DMC1}$*-GFP-lacI*], *can1-R,* $P_{CLB2}$*-3HA-SGO1 KANMX6, leu2::*OPL46[$P_{URA3}$*-tetR-tdTomato leu2::HIS3*] OPL214[*CEN3-5.1KB tetO256 URA3 TRP1 ARS1*] | *sgo1-md* | Fig. 5 A, B<br><br>Fig. 6 A, B |
| | Y2177: *MAT∝, trp1-Δ63, his3-Δ1, leu2, lys2::*pMDE798[$P_{DMC1}$*-GFP-lacI*], *tyr1-2, met13-c, cyh2-1,* $P_{CLB2}$*-3HA-SGO1 KANMX6,* OPL210[*CEN3-5.1KB lacO256 LEU2 TRP1 ARS1*] | | |
| DJE90 | X3104: *MAT∝, ura3-13, trp1-Δ63, leu2-?, tyr1-1, lys2-1, can1-R, his3::pHC30[pDMC1-3x yEVenus-lacI-I12 HIS3], SPC42-[MDE1145: URA3::HIS SPC42-DSRed], ura3::pKB80 [*$P_{GPD1}$*-GAL4(848)-ER-URA3::hphNT1], natNT2-*$P_{GAL1}$*-NDT80, met13-d, OPL210 [lacO, CEN3, TRP1/ARS1, LEU2]* | *SGO1* | Fig. 5 D |
| | Y2737: *MATa, leu2, lys2-2, met13-c, tyr1-2, ura3-1, trp1-Δ63, cyh2-1, his3::pHC29[*$P_{URA3}$*-tetR-3XmTurquoise-9MYC HIS3], trp1::pD280[*$P_{DMC1}$*-3XyEVenus-lacI-I12 TRP1], natNT2-*$P_{GAL1}$*-NDT80, OPL214 [tetO, CEN3, TRP1/ARS1, URA3]* | | |
| DJE91 | X3426: *MAT∝, ura3-13, trp1-Δ63, leu2-?, tyr1-1, lys2-1, can1-R, his3::pHC30[*$P_{DMC1}$*-3x yEVenus-lacI-I12 HIS3], SPC42-[MDE1145: URA3::HIS SPC42-DSRed], ura3::pKB80 [*$P_{GPD1}$*-GAL4(848)-ER-URA3::hphNT1], natNT2-*$P_{GAL1}$*-NDT80, met13-d, sgo1::KanMX4, OPL210 [lacO, CEN3, TRP1/ARS1, LEU2]* | *sgo1Δ* | Fig, 5 D |
| | Y2893: *leu2-?, lys2-2, met13-c, tyr1-2, MATa, ura3-1, trp1-Δ63, cyh2-1, his3::pHC29[pURA3-tetR-3XmTurquoise-9MYC HIS3], trp1::pD280[*$P_{DMC1}$*-3XyEVenus-lacI-I12 TRP1], natNT2-*$P_{GAL1}$*-NDT80, sgo1::KanMX4, OPL214 [tetO, CEN3, TRP1/ARS1, URA3]* | | |
| DJE99 | X3605: *MATa, ura3-13, trp1-Δ63, leu2-?, tyr1-1, lys2-1, met13-d, can1-R, his3::pHC30[*$P_{DMC1}$*-3x* | *sgo1-3A* | Fig. 6 B |

| | | | |
|---|---|---|---|
| | *yEVenus-lacI-I12 HIS3], SPC42-[MDE1145: URA3::HIS SPC42-DSRed], sgo1::sgo1-3a, OPL210 [lacO, CEN3, TRP1, LEU2]* | | |
| | Y2931: *MAT∝, leu2-?, lys2-2, met13-c, tyr1-2, ura3-1, trp1-Δ63, cyh2-1, his3::pHC29[$P_{URA3}$-tetR-3XmTurquoise-9MYC HIS3], trp1::pD280[$P_{DMC1}$-3XyEVenus-lacI-I12 TRP1], sgo1::sgo1-3a, OPL214 [tetO, CEN3, TRP1/ARS1, URA3]* | | |
| DEK314 | X2333: *MAT∝, ura3-13, trp1-63, leu2, leu2::*OPL46[*$P_{URA3}$-tetR-tdTomato leu2::HIS3*], *met13-d, tyr1-1, can1-R, lys2::*pMDE798[*$P_{DMC1}$-GFP-lacI*], OPL210[*CEN3-5.1KB, lacO256, LEU2 TRP1, ARS1*], OPL214[*CEN3-5.1KB tetO256 URA3 TRP1 ARS1*] | *SGO1* | Fig. 6 B |
| | Y2029: *MAT***a**, trp1-Δ63, his3-Δ1, leu2, lys2::pMDE798[$P_{DMC1}$-GFP-lacI], tyr1-2, met13-c, cyh2-1 | | |
| DEK305 | X2349: MAT**a**, *trp1-Δ63, his3-Δ1, leu2, met13-d, tyr1-1, lys2::*pMDE798[*$P_{DMC1}$-GFP-lacI*], *can1-R, $P_{CLB2}$-3HA-SGO1 KANMX6, leu2::*OPL46[$P_{URA3}$-*tetR-tdTomato leu2::HIS3*], OPL210[*CEN3-5.1KB lacO256 LEU2 TRP1 ARS1*] | *sgo1-md* | Fig. 6 B |
| | Y2175: *MAT∝, trp1-Δ63, his3-Δ1, leu2, lys2::*pMDE798[*$P_{DMC1}$-GFP-lacI*], *tyr1-2, met13-c, cyh2-1, $P_{CLB2}$-3HA-SGO1 KANMX6,* OPL214[*CEN3-5.1KB tetO256 URA3 TRP1 ARS1*] | | |
| DMS223 | DMS181.9C: *MAT∝, his3-11,15, arg4, ade1::ARG4, cup1::ura3::THR, trp2, leu2 S. cerevisiae* chromosome *V*: r*ad3, ilv1, ura3::HIS3::*AFS152[*URA3 $P_{CYC1}$-GFP-lacI*], *sec3::* pBK13.1[*LEU2 lacO*] | *SGO1* Exchange | Fig. 6 C |
| | DMS181.3B: *MAT***a**, *his3-11,15, arg4, ade1::ARG4, cup1::ura3::THR, trp2, lys1, leu2 S. cerevisiae* chromosome V: *rad3, ilv1, ura3::HIS3::* pAFS152[*URA3 $P_{CYC1}$-GFP-lacI*], *sec3::*pBK13.1[*LEU2 lacO*] | | |
| DMS218 | DMS202.10D: *MAT***a**, *ade1::ARG4, cup1::ura3::THR4, leu2, his3, arg4, lys1, sgo1::KANMX6 S. cerevisiae* chromosome *V*: *rad3, ilv1, ura3::HIS3::*pAFS152[*URA3 $Pr_{CYC}$-GFP-lacI*], *sec3::*pBK13.1[*LEU2 lacO*] | *sgo1Δ* Exchange | Fig. 6 C |
| | DMS202.1A: *MAT∝, ade1::ARG4, cup1::ura3::THR4, trp1, leu2, his3, arg4, lys1, sgo1::KANMX6, S. cerevisiae* chromosome *V*: *rad3, ilv1, ura3::HIS3:: pAFS152*[*URA3 $P_{CYC}$-GFP-lacI*], *sec3::pBK13.1*[*LEU2 lacO*] | | |
| DMS181 | TMS46-7.3D: *MATα, ade1::ARG4, trp2, leu2,his3-11,15, arg4ΔHpa, cup1::ura2::THR4,* | *SGO1* | Fig. 6 D |

| | | | |
|---|---|---|---|
| | *lys1, S. cerevisiae* chromosome *V: rad3, ilv1, ura3::HIS3:: pAFS152*[*URA3 $P_{CYC}$-GFP-lacI*], *sec3::*pBK13.1[*LEU2-lacO*]<br>DMS143.16A: *MAT***a**, *his3-11,15, lys2-801::*pLL1.1[*LYS2 $P_{CYC}$GFP lacI*], *arg4ΔHpa, cup1::ura3::THR1*<br>*S. bayanus* chromosome *V: URA3, pac2::* pD174*[LEU2 lacO], ilv1* | Non-exchange | |
| DMS217 | DMS202.1A: *MAT∝,ade1::ARG4, cup1::ura3::THR, trp1, leu2, his3, arg4, lys1, sgo1::KANMX6 S. cerevisiae* chromosome *V*: *rad3, ilv1, ura3::HIS3::*pAFS152*[URA3 $P_{CYC}$-GFP-lacI], sec3::*pBK13.1[*LEU2 lacO*]<br>DMS207.1A: *MAT***a**, *his3, lys2-801::* pLL1.1[ *LYS2 $P_{CYC}$GFP lacI*], *arg4, cup1::ura3::THR1, spo11::natNT2 S. bayanus* chromosome *V: URA3, pac2::*pD174[*LEU2 lacO*], *ilv1* | *sgo1Δ*<br>Non-exchange | Fig. 6 D |
